# Dietary intake and diet quality of Swiss adult survivors of childhood cancer compared to the general population

**DOI:** 10.1101/527424

**Authors:** Fabiën N. Belle, Angeline Chatelan, Rahel Kasteler, Idris Guessous, Maja Beck Popovic, Marc Ansari, Claudia E. Kuehni, Murielle Bochud, for the Swiss Pediatric Oncology Group (SPOG)

## Abstract

**Background:** Childhood cancer survivors (CCSs) are at increased risk of developing chronic health conditions, which may be reduced by following a healthy lifestyle including a good diet.

**Objectives:** This study compared the dietary intake and quality of CCSs and the general population.

**Design:** As part of the Swiss Childhood Cancer Survivor Study, we sent a food frequency questionnaire (FFQ) to CCSs who had a median age of 34 years (IQR: 29-40 years) and a median of 26 years (20-31 years) postdiagnosis at the time of survey. We compared dietary intake and quality of CCSs and three comparison groups representing the general adult population using FFQ and 24h recall data (24HDR). We evaluated whether mean individual intake met national dietary recommendations and used the Alternative Healthy Eating Index (AHEI) to estimate diet quality.

**Results:** The 774 CCSs in our study were compared to 8964 participants in the Bus Santé study, 1276 participants in the CoLaus study, and 1134 participants in the Swiss National Nutrition Survey. Dietary intake was equally poor in CCSs and the general Swiss population. CCSs consumed inadequate amounts of vitamin D, fiber, carbohydrates, iron, vitamin A, and calcium (12%, 41%, 72%, 72%, 79%, and 89% of the recommended intakes, respectively), and excessive amounts of saturated fat, protein, cholesterol, and total fat (137%, 126%, 114%, and 107% of the recommended intakes). The mean AHEI score in CCSs was low at 48.0 (men: 45.0, women: 50.9) out of a maximum score of 100. The general population, assessed by 24HDR, scored lower overall than CCSs (41.5; men: 38.7, women: 43.8). Clinical characteristics were not associated with diet quality in CCSs.

**Conclusion:** Long-term CCSs and the general adult population have similarly poor dietary intake and quality in Switzerland, which suggests population-based interventions for everyone.

## INTRODUCTION

A healthy diet is a modifiable factor that can prevent or delay the development of chronic diseases such as type II diabetes, metabolic syndrome, and cardiovascular disease (CVD) (1–3). Populations suffering from these comorbidities are widely recommended to consume a diet rich in fruit, vegetables, fiber, and complex carbohydrates, which is low in saturated and trans-fatty acids, and to consume alcohol in moderation (1–3). Accumulating research among childhood cancer survivors (CCSs) shows that the burden of chronic diseases secondary to childhood cancer or its treatment can be reduced with dietary adaptation, weight management, and physical activity (4–7). A healthy diet is therefore particularly relevant for CCSs who are at increased risk of developing chronic diseases, yet few studies have evaluated the intake or quality of the diet of CCSs (8, 9).

Poor adherence to the 2010 Dietary Guidelines for Americans and poor diet quality based on the Healthy Eating Index-2010 (HEI-2010, 58% of the maximum score) were found in 2570 adult long-term CCSs in the St. Jude Lifetime cohort (10). Diet quality was even poorer in those CCSs diagnosed young and treated with abdominal radiation therapy. Similar results have been observed in other, smaller studies in the United States (US): CCSs poorly adhered to dietary recommendations and mean diet quality ranged from 33% to 56% of the maximum score (5, 6, 11–13).

Studies involving different lifestyles and eating habits outside the US are few and they have small sample sizes and short follow-up times, or focus on specific cancer diagnoses (14, 15). Almost all studies performed to date lack control groups to compare dietary intake and quality between CCSs and the general population. Therefore, we analyzed data from the Swiss Childhood Cancer Survivor Study (SCCSS) to assess dietary intake and diet quality of CCSs and compared them to those of the general Swiss adult population. The comparison with the general population is important to see whether dietary intake and quality are influenced by cancer and/or its treatment. We also explored whether selected baseline clinical characteristics in CCSs were associated with dietary intake at follow-up.

## METHODS

### Study populations

#### The Swiss Childhood Cancer Survivor Study (SCCSS)

The SCCSS is a population-based, long-term follow-up study of all childhood cancer patients registered in the Swiss Childhood Cancer Registry (SCCR; available from: www.childhoodcancerregistry.ch) who were diagnosed in Switzerland with leukemia, lymphoma, central nervous system (CNS) tumors, malignant solid tumors, or Langerhans cell histiocytosis; who were under the age of 21 years at time of diagnosis; who survived ≥5 years after initial diagnosis of cancer; and who were alive at the time of the study (16–18). Ethical approval of the SCCR and the SCCSS was granted by the Ethics Committee of the Canton of Bern (KEK-BE: 166/2014). This study is registered at clinicaltrials.gov as NCT03297034. As part of the SCCSS, we traced all addresses of CCSs diagnosed between 1976 and 2005 who had filled in the baseline questionnaire between 2007 and 2013 (N=2527, response rate 70%) (16) and then sent a follow-up questionnaire in February 2017. Nonresponders received a reminder about the follow-up questionnaire eight weeks later. If they again did not respond, we sent them a second reminder. Our questionnaire included core questions from the US and UK CCS studies (19, 20), with added questions about dietary intake (21, 22), health behaviors, and sociodemographic measures from the Swiss Health Survey (SHS) (23) and the Swiss Census (24).

#### Comparison groups

We used three random samples of the general Swiss adult population represented by data from the Bus Santé survey, the Cohorte Lausannoise (CoLaus), and the Swiss National Nutrition Survey (menuCH). Bus Santé is a cross-sectional, population-based survey that is ongoing in the canton of Geneva (25). A representative sample of non-institutionalized men and women aged 35-74 years has been recruited each year since 1993. Eligible participants are identified with a standardized procedure using a residential list established by the local government. Random sampling in age and sex-specific strata is proportional to the corresponding frequencies in the population. Those who did not respond after three mailings and seven phone calls were replaced using the same selection protocol, but those who declined to participate were not replaced. Participants were not eligible for future recruitments and surveys. Participants received a self-administered questionnaire including a food frequency questionnaire (FFQ, described below) to collect data on sociodemographic characteristics, health behaviors, and dietary intake at home before receiving an invitation to a health examination in a clinic or a mobile medical unit. During the examination, trained staff checked the questionnaires for completeness.

The CoLaus study (www.colaus-psycolaus.ch) is a prospective, population-based cohort study conducted in the city of Lausanne to identify biological and genetic determinants of CVD. From June 2003 until May 2006, participants aged 40-75 years were recruited for the baseline examination. Those who participated in the baseline study were asked to participate in the follow-up study between April 2009 and September 2012 (26). In the follow-up study, dietary intake was assessed with the same FFQ that was used in the Bus Santé surveys.

The Swiss National Nutrition Survey (www.menuch.ch) is a cross-sectional nutrition survey conducted from January 2014 to February 2015 among persons 18 to 75 years old in the three main linguistic regions (German, French, and Italian) of Switzerland (22). Trained dieticians collected the data. Participants answered two 24-hour dietary recalls (24HDR), the first face-to-face and the second by phone two to six weeks later. Prior to the face-to-face interview, participants received a 49-item questionnaire including questions on sociodemographic characteristics and health behaviors. Interviews were carried out in German, French, and Italian.

### Measurements

#### Dietary intake

Dietary intake of CCSs was assessed in participants in the Bus Santé and CoLaus surveys using the same self-administered, semiquantitative FFQ including portion sizes (27, 28). The FFQ was originally developed and validated for the French-speaking Swiss adult population (21, 25, 27, 29). It collects information on consumption frequency and portion sizes of 97 fresh and prepared food items (not including dietary supplements), organized in 12 different food groups, during the previous four weeks. Consumption frequencies range from never during the last four weeks to two or more times per day, and portions are divided into three sizes equal to, or smaller or larger than a reference size. The reference portions are defined as common household measures representing the median portion size of a previous validation study performed with 24HDRs (21). The smaller and larger portion sizes represented the first and third quartiles of this distribution. The French Information Center on Food Quality (Centre d’Information sur la Qualité des Aliments) food composition table was used to convert the food portions into macronutrients and micronutrients.

In the Swiss National Nutrition Survey dietary intake was assessed by two nonconsecutive 24HDRs covering all seasons and weekdays, including weekends, by using GloboDiet software (30, 31). Each 24HDR was complemented with a comprehensive picture book (32) and a set of real dishes adapted to the Swiss-specific food market to support survey participants in quantifying amounts of foods consumed. Conversion into macro- and micronutrients was performed using the Swiss Food Composition Database (SFCDB) (33). We averaged the dietary intake of the two 24HDRs for each participant.

Because the original FFQ was developed for the French-speaking part of Switzerland, we extended the FFQ with 15 additional food items for CCSs based on data from the Swiss National Nutrition Survey (22). We investigated whether foods most frequently consumed in the German and Italian-speaking part of Switzerland were also included in the FFQ. The 15 additional food items were reported more than 70 times (>2% of the total food items) during the 24h recall assessments.

#### Sociodemographic, lifestyle, and clinical characteristics

For all CCSs and comparison groups, we collected self-reported data on sex, age at survey, educational level, country of birth, language region in Switzerland, household status, physical activity, smoking status, and BMI at survey. Physical activity was assessed differently in CCSs and comparison groups. In the SCCSS and the Swiss National Nutrition Survey, we dichotomized physical activity into two groups according the WHO guidelines for physical activity in adults: inactive (lower than 150 minutes of activity per week); and active (150 minutes or more of moderate 75 minutes of vigorous physical activity, or a combination of moderate and vigorous intense physical activity per week) (34). In the Bus Santé and CoLaus studies, physical activity was assessed with a validated, self-administered physical activity frequency questionnaire (PAFQ) (35) and dichotomized into lower (inactive), or equal or more than the first quartile of total weekly physical activity time excluding sleep (active). For all CCSs, we had information on weight without clothes and height without shoes at time of survey from the self-administered questionnaires. In the three random samples of the general Swiss adult population, participant body weight and height were measured without shoes, in light indoor clothes. Body weight was measured in kilograms to the nearest 100 g using a calibrated electronic scale (Seca^®^, Hamburg Germany). Height was measured to either the nearest 5 or 10 mm using a Seca^®^ height gauge. We calculated BMI by dividing weight in kilograms by height in meters squared (kg/m^2^) in all groups. BMI in adults was classified as underweight (<18.5 kg/m^2^), normal (≥18.5 to <25 kg/m^2^), overweight (≥25 to <30 kg/m^2^), or obese (≥30 kg/m^2^) (36). For the CCSs population, we extracted additional clinical information from the Swiss Childhood Cancer Registry (SCCR). This included information on cancer diagnosis and the age at diagnosis. Diagnosis was classified according to the International Classification of Childhood Cancer, 3^rd^ Edition (37). Radiotherapy was classified as any, cranial, chest, total body and/or abdominal, or no radiotherapy. Cranial radiation was considered as present if the survivor had received direct radiation to the brain and/or skull. Chest radiotherapy was regarded as direct radiation applied to the chest including total body irradiation, mantlefield irradiation, or irradiation of the thorax, mediastinum, or thoracic spine. Cumulative dosage of radiotherapy was obtained from medical records, and categorization was based on the Children’s Oncology Group Long-Term Follow-up (COG-LTFU) Guidelines. Irradiation was categorized as <18 Gray (Gy) or ≥18 Gy for cranial irradiation; the chest, <30 Gy or ≥30 Gy; and as total body and/or abdominal irradiation versus no radiation (38). Other treatment exposures were divided into glucocorticoids, anthracyclines, alkylating agents, and hematopoietic stem cell transplantation (HSCT). Glucocorticoids intake, including prednisone and/or dexamethasone, was based on cancer protocol adherence as described previously (39). We also retrieved records on relapse during follow-up.

### Statistical Analyses

We included all CCSs and participants from the general population (in the Bus Santé, CoLaus, Swiss National Nutrition surveys) who were aged 20-50 years at time of survey, who provided reliable dietary intake information, and were neither pregnant nor lactating during the survey (**Supplemental Figure 1**). For better comparison between CCSs and peers, we standardized comparison groups for sex, age at survey, and language region when applicable as previously described (40–42). The first step in our analyses was to evaluate whether CCS and their peers met the dietary recommendations for Germany (D), Austria (A), and Switzerland (CH) (DACH) (43). We compared mean intake to the recommended intake or, when not available, the adequate intake. We calculated mean intake based on age and sex recommendations weighted by the age and sex distribution of the study population. Nutritional goals were set at 100 when the mean intake met the recommended or adequate intake. Total energy intake was calculated including calories from alcohol consumption. We used the same FFQ with CCSs and Bus Santé and CoLaus participants for direct comparability. To estimate the diet quality of CCSs and peers we used the Alternative Healthy Eating Index (AHEI) (44). All AHEI score components range from zero (worst) to ten (best), and the total AHEI score ranges from zero (nonadherence) to 100 (perfect adherence). To calculate the AHEI, we used the extended FFQ version with the 15 additional food products based on data of the Swiss National Nutrition Survey. We compared whether the AHEI score differed by sex between CCSs and the Swiss National Nutrition Survey. We assessed how dietary quality of CCSs differed by cancer diagnosis and treatment with ANCOVA while adjusting for sex, age, and cancer diagnosis. We did not adjust for education level, smoking habits, physical activity, and BMI because these covariates can be affected by cancer diagnosis, treatment exposures, and the occurrence of late effects (i.e., intermediates on the causal pathway). We used Stata (version 14, Stata Corporation, Austin, Texas) for all analyses.

## RESULTS

### Response rate and characteristics of the study populations

Among 1749 eligible CCSs, we traced and contacted 1607, 918 of whom (57%) returned a questionnaire. We excluded 29 survivors who were over 50 years old, 11 who were pregnant or lactating, 34 who did not report their dietary intake, and a further 70 who had unreliable dietary intake data <850 kcal or >4500 kcal per day. We thus included 774 CCSs in this study (**Supplemental Figure S1**). Participation rates in the Bus Santé survey ranged from 55 to 75% (45). Of 20,125 participants in the Bus Santé survey, 10,851 were over 50 years old, and 310 had unreliable dietary intake information. We thus included 8964 Bus Santé participants in this study. Of 5064 participants in the CoLaus study representing a participation rate of 41%, 3616 were excluded because they were over 50 years old, 126 because they had no dietary intake information, and 46 because of unreliable dietary intake, leaving 1276 participants for the analyses. In the Swiss National Nutrition Survey the participation rate was 38%. From 2085 participants (38% response rate) we excluded 857 who were older than 50 years 39 who were younger than 20 years, 27 who were pregnant or lactating, one participant who did not report his dietary intake, and 27 participants with unreliable dietary intake information, leaving 1134 participants for the analyses.

More CCSs than controls were born in Switzerland since the initial inclusion criterion of the SCCR stipulated living in Switzerland at time of diagnosis (all *P* < 0.001). CCSs earned a university degree less often than those in the Bus Santé and menuCH studies (all *P* < 0.001). CCSs smoked less (all *P* < 0.001) and more often had a normal BMI (*P*_Bus Santé_ < 0.001, *P*_CoLaus_ < 0.001, *P*_menuCH_ = 0.023) than controls. Characteristics of CCSs and control are given in **Table I**.

**Table I.** Characteristics of childhood cancer survivors and comparison groups^1^

| Characteristics | CCSs<br>(n = 774) | Bus Santé<sup>2</sup><br>(n = 8964) |  | CoLaus<sup>2</sup><br>(n = 1276) |  | menuCH<sup>2</sup><br>(n = 1134) |  |
| --- | --- | --- | --- | --- | --- | --- | --- |
|  |  | n (%<sub>std</sub>) | P<sup>3</sup> | n (%<sub>std</sub>) | P<sup>3</sup> | n (%<sub>std</sub>) | P<sup>3</sup> |
| **Sex, n (%)** |  |  |  |  |  |  |  |
| Men | 386 (50) | 4345 (55) | NA | 628 (49) | NA | 498 (51) | NA |
| **Age at survey, n (%)** |  |  |  |  |  |  |  |
| 20-24 yrs | 69 (9) | - | NA | - | NA | 169 (9) | NA |
| 25-29 yrs | 179 (23) | - |  | - |  | 188 (23) |  |
| 30-34 yrs | 163 (21) | - |  | - |  | 150 (22) |  |
| 35-39 yrs | 156 (20) | 3027 (45) |  | - |  | 146 (20) |  |
| 40-44 yrs | 123 (16) | 2997 (32) |  | 552 (62) |  | 211 (16) |  |
| 45-50 yrs | 84 (11) | 2940 (23) |  | 724 (38) |  | 270 (11) |  |
| **Language region, n (%)** |  |  |  |  |  |  |  |
| German speaking | 554 (72) | - | NA | - | NA | 740 (71) | NA |
| French speaking | 202 (26) | 8964 (100) |  | 1276 (100) |  | 280 (27) |  |
| Italian speaking | 18 (2) | - |  | - |  | 114 (2) |  |
| **Country of birth, n (%)** |  |  |  |  |  |  |  |
| Switzerland | 737 (95) | 4811 (54) | <0.001 | 750 (58) | <0.001 | 934 (81) | <0.001 |
| Other | 37 (5) | 4153 (46) |  | 526 (42) |  | 200 (19) |  |
| **Education (highest degree), n (%)** |  |  |  |  |  |  |  |
| Lower than university | 537 (69) | 5336 (59) | <0.001 | 916 (71) | 0.817 | 741 (61) | <0.001 |
| University | 237 (31) | 3628 (41) |  | 360 (29) |  | 393 (39) |  |
| **Household status, n (%)** |  |  |  |  |  |  |  |
| Alone | 156 (20) | 2192 (24) | 0.013 | 237 (20) | 0.765 | 170 (16) | 0.029 |
| Other | 618 (80) | 6772 (76) |  | 1039 (80) |  | 964 (84) |  |
| **Physical activity<sup>4</sup>, n (%)** |  |  |  |  |  |  |  |
| Inactive | 159 (21) | 662 (8) | <0.001 | 267 (21) | <0.001 | 220 (19) | 0.051 |
| Active | 613 (79) | 2700 (33) |  | 905 (71) |  | 900 (80) |  |
| Missing | 2 (<1) | 5602 (58) |  | 104 (8) |  | 14 (1) |  |
| **Smoking status, n (%)** |  |  |  |  |  |  |  |
| Never | 523 (68) | 4005 (44) | <0.001 | 567 (46) | <0.001 | 521 (44) | <0.001 |
| Former | 129 (17) | 2460 (27) |  | 398 (30) |  | 320 (28) |  |
| Current | 122 (16) | 2499 (29) |  | 311 (24) |  | 293 (28) |  |
| **BMI at survey, n (%)** |  |  |  |  |  |  |  |
| Underweight | 39 (5)<sup>5</sup> | 241 (3) | <0.001 | 19 (1) | <0.001 | 34 (3) | 0.024 |
| Normal | 491 (63) | 5536 (62) |  | 649 (49) |  | 687 (60) |  |
| Overweight | 170 (22) | 2491 (28) |  | 451 (38) |  | 306 (28) |  |
| Obese | 74 (10) | 696 (8) |  | 157 (12) |  | 107 (9) |  |

<sup>1</sup> BMI, body mass index; CCS, childhood cancer survivors; NA, not applicable; std, standardized.<sup>2</sup> Standardized on sex, age at survey, and language region according to CCSs.<sup>3</sup> P value calculated from chi-square statistics comparing control groups with CCSs (2-sided test).<sup>4</sup> CCSs and Swiss National Nutrition Survey (menuCH): < or ≥150 minutes of moderate, or 75 minutes of vigorous physical activity or a combination of moderate and vigorous activity per week. Bus Santé and CoLaus: < or ≥ 1<sup>st</sup> quartile of total weekly physical activity time excluding sleep.<sup>5</sup> Self-reported BMI in CCSs.

Among CCSs, the most common cancers were leukemia, lymphoma, and CNS tumors (**Supplemental Table S1**). Median age at diagnosis was nine years (IQR: 4–14 years) and median time from diagnosis to survey was 26 years (IQR: 20–31 years). Eleven percent had experienced a relapse. CCSs got treatment exposure to glucocorticoids (43%), alkylating agents (41%), anthracyclines (38%), radiation (36%), and hematopoietic stem cell transplantation (4%).

### Dietary intake and diet quality in CCSs and comparison groups

Reported total energy intake was low across all studies (**Table II**), in particular in CCSs. Participants in menuCH reported higher energy intake, assessed by 24HDRs, than participants in the other studies for whom the assessment method was FFQ. Dietary intakes of CCSs and comparison groups compared to the dietary DACH recommendations are represented in **Figure 1**. Although the differences in percentages of the recommended DACH intake or limit were statistically significant between CCSs and the comparison groups, the absolute differences were small and potentially the result of different dietary assessment tools (**Table II** and **Figure 1**). Nevertheless, CCSs had a substantially lower daily intake of alcohol (5.7g vs. Bus Santé: 11.7g, CoLaus: 8.7g, and menuCH: 11.8g). In CCSs, intakes higher than recommended, considering age and sex distributions, were seen for saturated fat (137% of the recommended intake), protein (126%), cholesterol (114%), and total fat (107%). Intakes lower than recommended were seen for vitamin D (12%), fiber (41%), carbohydrates (72%), iron (72%), vitamin A (79%), and calcium (89%). We found no large difference in adherence to DACH recommendations between sexes in CCSs, except for cholesterol (men: 125%, women: 103%) and iron (men: 97%, women: 56%) (**Supplemental Table S2**).

**Figure 1.**
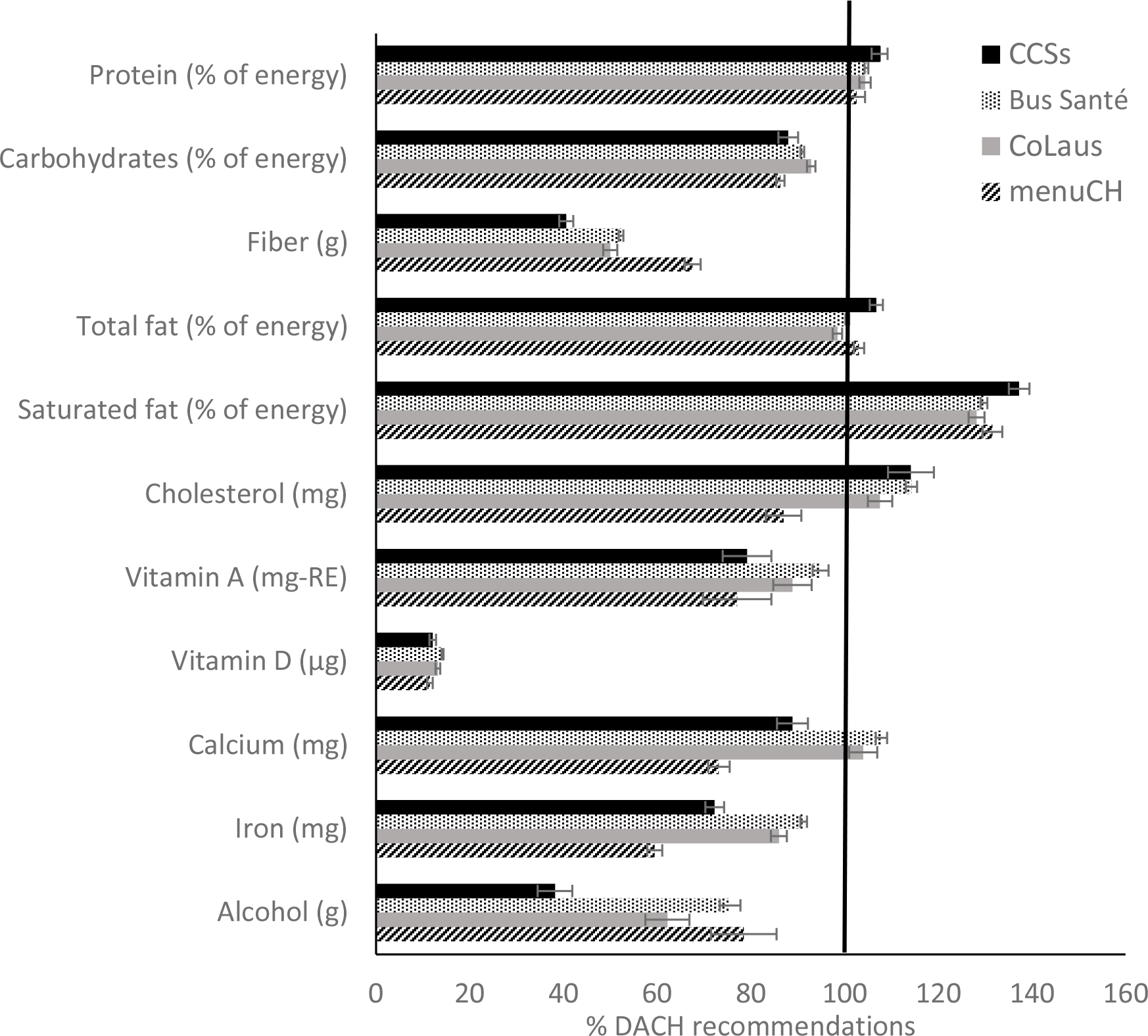
Nutrient intake compared with DACH recommended intake or limit in childhood cancer survivors and the general Swiss population1: Bus Santé, CoLaus, and menuCH^1^. The length of the bar per nutrient corresponds to the percentage of mean intake (95% CIs) compared to the recommended intake level * 100. Recommended intake is estimated on the basis of age and sex according to dietary recommendations for Germany (D), Austria (A) and Switzerland (CH) (DACH) 2015, weighted by the age and sex distribution per study population. For alcohol intake the maximum tolerated dosage was taken. Nutritional goals were set at 100 when the mean intake met the recommended intake or limit. ^1^ Standardized on sex, age at survey, and language region according to CCSs.

**Table II.** Dietary intake in childhood cancer survivors and the general Swiss population: Bus Santé, CoLaus, and menuCH compared to the DACH dietary recommendations^1^

| Nutrients | DACH recommendations<sup>3</sup> |  | CCSs (n = 774) |  | Bus Santé<sup>2</sup> (n = 8964) |  | CoLaus<sup>2</sup> (n = 1276) |  | menuCH<sup>2</sup> (n = 1134) |  |
| --- | --- | --- | --- | --- | --- | --- | --- | --- | --- | --- |
|  | men | women | mean ± SD | % DACH<sup>4</sup> | mean ± SD | % DACH<sup>4</sup> | mean ± SD | % DACH<sup>4</sup> | mean ± SD | % DACH<sup>4</sup> |
| Total energy, kcal<sup>5</sup> | 2300-3400 | 1800-2600 | 1639 ± 582 | 65 | 2022 ± 688 | 79 | 1923 ± 668 | 76 | 2283 ± 720 | 90 |
| **Macronutrients** |  |  |  |  |  |  |  |  |  |  |
| Protein, g | 57 | 48 | 66.1 ± 29.5 | 126 | 78.7 ± 29.5 | 149 | 74.8 ± 29.4 | 143 | 86.9 ± 37.2 | 165 |
| % of energy<sup>6</sup> | 10-20 | 10-20 | 16.1 ± 3.6 | 108 | 15.7 ± 3.1 | 105 | 15.7 ± 3.3 | 104 | 15.4 ± 4.6 | 103 |
| Vegetal, g |  |  | 18.1 ± 7.6 |  | 23.6 ± 10.0 |  | 22.2 ± 9.6 |  |  |  |
| Animal, g |  |  | 48.0 ± 26.9 |  | 55.1 ± 24.8 |  | 52.6 ± 24.4 |  |  |  |
| Carbohydrates, g<sup>7</sup> | 225-275 | 225-275 | 180 ± 74 | 72 | 230 ± 91 | 92 | 224 ± 89.9 | 90 | 246 ± 89 | 98 |
| % of energy<sup>6,7</sup> | 45-55 | 45-55 | 43.9 ± 8.8 | 88 | 45.5 ± 8.6 | 91 | 46.5 ± 8.1 | 93 | 43.2 ± 8.1 | 86 |
| Mono/disaccharides, g |  |  | 86 ± 46 |  | 105 ± 52 |  | 107 ± 53 |  | 110 ± 54 |  |
| Polysaccharides, g |  |  | 94 ± 44 |  | 124 ± 60 |  | 116 ± 57 |  | 83 ± 49 |  |
| Total fiber, g<sup>7</sup> | ≥30 | ≥30 | 12.2 ± 6.4 | 41 | 15.7 ± 8.0 | 52 | 15.0 ± 8.0 | 50 | 20.3 ± 8.9 | 68 |
| Total fat, g |  |  | 68.2 ± 28.2 |  | 79.0 ± 31.1 |  | 73.5 ± 28.7 |  | 91.8 ± 35.1 |  |
| Total fat, % of energy<sup>6</sup> | 30-40 | 30-40 | 37.4 ± 6.9 | 107 | 35.2 ± 6.8 | 101 | 34.5 ± 6.2 | 99 | 36.1 ± 6.9 | 103 |
| Saturated fat, g |  |  | 25.2 ± 11.6 |  | 29.4 ± 13.4 |  | 27.5 ± 12.4 |  | 33.6 ± 14.8 |  |
| Saturated fat, % of energy<sup>8</sup> | <10 | <10 | 13.7 ± 3.2 | 137 | 13.0 ± 3.3 | 130 | 12.8 ± 3.1 | 128 | 13.2 ± 3.6 | 132 |
| Monounsaturated fat, g |  |  | 28.6 ± 12.7 |  | 31.4 ± 13.0 |  | 29.4 ± 11.9 |  | 23.2 ± 12.6 |  |
| Polyunsaturated fat, g |  |  | 8.6 ± 3.8 |  | 11.6 ± 5.3 |  | 10.3 ± 4.7 |  | 8.6 ± 5.3 |  |
| Cholesterol, mg<sup>6</sup> | 300 | 300 | 343 ± 208 | 114 | 343 ± 167 | 114 | 323 ± 141 | 108 | 261 ± 194 | 87 |
| **Micronutrients** |  |  |  |  |  |  |  |  |  |  |
| **Vitamins** |  |  |  |  |  |  |  |  |  |  |
| A, mg-RE<sup>9</sup> | 1.0 | 0.8 | 0.7 ± 0.7 | 79 | 0.9 ± 0.7 | 99 | 0.8 ± 0.7 | 89 | 0.7 ± 1.1 | 77 |
| D, µg | 20 | 20 | 2.4 ± 2.0 | 12 | 2.8 ± 2.1 | 14 | 2.6 ± 1.9 | 13 | 2.3 ± 2.2 | 11 |
| **Minerals** |  |  |  |  |  |  |  |  |  |  |
| Calcium, mg | 1000 | 1000 | 889 ± 471 | 89 | 1079 ± 563 | 108 | 1040 ± 543 | 104 | 732 ± 391 | 73 |
| Iron, mg | 10<sup>4</sup> | 15 | 9.0 ± 3.6 | 72 | 11.2 ± 3.9 | 91 | 10.8 ± 3.9 | 86 | 7.4 ± 3.5 | 59 |
| Alcohol, g<sup>10</sup> | 20 | 10 | 5.7 ± 7.8 | 38 | 11.7 ± 16.4 | 75 | 9.3 ± 12.6 | 62 | 11.8 ± 18.0 | 78 |

<sup>1</sup> CCS, childhood cancer survivors, DACH, Dietary recommendations for Germany (D), Austria (A) and Switzerland (CH).
<sup>2</sup> Standardized on sex, age at survey, and language region according to CCSs.
<sup>3</sup> DACH recommendations for the general population age 20-50 years, excluding pregnant and lactating women, 2015.
<sup>4</sup> Percentage of mean intake in relation to the DACH recommended intake level \* 100. Recommended intake is estimated on the basis of the age-sex groups of the DACH guidelines, weighted by the age and sex distribution of the study population. For alcohol intake the maximum tolerated dosage was taken.
<sup>5</sup> Depending on physical activity level (PAL, 1.4-1.8) and age.
<sup>6</sup> Federal Food Safety and Veterinary Office FSVO, www.naehrwertdaten.ch.
<sup>7</sup> BLV (2009). Kohlenhydrate in unserer Ernährung – Empfehlungen des BLV. www.blv.admin.ch.
<sup>8</sup> BLV (2012). Fett in unserer Ernährung – Empfehlung des BLV. www.blv.admin.ch, recommendations as a percentage of the daily energy demand and based on a daily energy intake of 2000 kcal.
<sup>9</sup> Total vitamin A= retinol + carotene/12, expressed in µg RAE.
<sup>10</sup> Maximum tolerated dosage.

The mean AHEI score in CCSs was low at 48.0 (men: 45.0, women: 50.9) out of a maximum score of 100, but it was higher than that observed in the general population (menuCH) (**Supplemental Table S3**). In CCSs, most of the individual components had a score of less than 50% of the maximum score (**Figure 2**). However, men’s scores were 52% for fish, 54% for alcohol, and 96% for trans fat (96%), while women scored 52% for polyunsaturated fatty acids (PUFA), 55% for vegetables, 58% for red and processed meat, and 95% for trans fat. MenuCH participant scores were lower overall than those of CCSs: 38.7 for men and 43.8 for women, and all individual components had a component score of less than 50% of the maximum score. Exceptions were PUFA, at 95% for men and 94% for women, and red and processed meat, which was 52% among women. The difference between CCSs and the general population was particularly large for vegetables, with higher scores for CCSs, whereas scores for sugar-sweetened beverages, alcohol, and nuts, seeds, legumes, and tofu were similar. Scores for red and processed meat were higher in CCSs than in menuCH participants.

**Figure 2.**
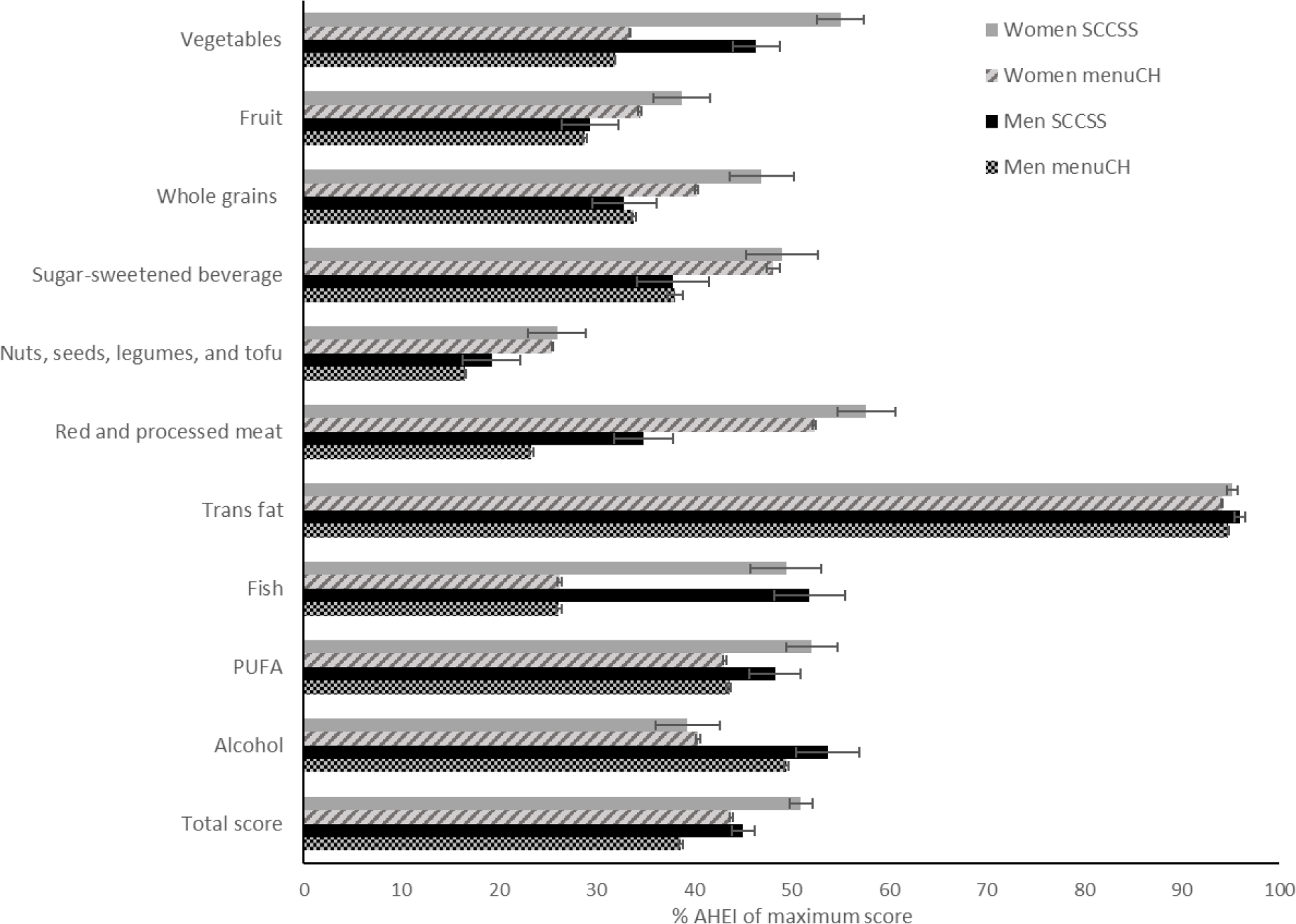
Alternate Healthy Eating Index^1,2^ scores with 95% CIs in childhood cancer survivors and the general population (menuCH)^3^ by sex. ^1^ Adapted from Chiuve et al. J Nutr 2012 142(6):1009-18. ^2^ Intermediate food intake was scored proportionately between the minimum score zero and the maximum score ten. ^3^ Adjusted for sex, age at survey, and if applicable ICCC-3 diagnosis, menuCH population additionally standardized on sex, age at survey, and language region according to CCSs.

### Clinical characteristics and dietary quality in CCSs

There was weak evidence that the dietary quality differed between types of cancer diagnosis (*P* = 0.078). CCSs of CNS tumors had an AHEI score of 45.5, whereas survivors of leukemia, lymphoma, and other cancers scored slightly higher (**Table III**). Among CCSs with CNS tumors, survivors of the largest subgroup (41%), astrocytomas, had an AHEI score of 44.1 (95% CI: 39.9, 48.3). Age at diagnosis, time since diagnosis, and history of relapse were not associated with diet quality in CCSs. Cancer treatment including cranial, chest, total body, and/or abdominal radiation; glucocorticoids, anthracyclines, alkylating agents, and hematopoietic stem cell transplantation were also not associated with diet quality.

**Table III.** Diet quality in childhood cancer survivors by clinical characteristics (retrieved from ANCOVA)^1, 2^

| Characteristics | CCSs<br>(n = 774) |  |  |
| --- | --- | --- | --- |
|  | n (%) | AHEI score<br>(95% CI) | P |
| **ICCC-3 diagnosis** |  |  |  |
| I: Leukemia | 237 (31) | 48.3 (46.8, 49.8) | 0.078 |
| II: Lymphoma | 163 (21) | 49.6 (47.8, 51.4) |  |
| III: CNS tumor | 79 (10) | 45.5 (42.9, 48.2) |  |
| Other | 295 (38) | 47.5 (46.2, 48.9) |  |
| **Age at diagnosis, yrs** |  |  |  |
| <5 | 251 (32) | 47.2 (45.5, 48.8) | 0.390 |
| 5-9 | 163 (21) | 48.7 (46.9, 50.6) |  |
| 10-14 | 224 (29) | 47.6 (46.0, 49.2) |  |
| 15-20 | 136 (18) | 49.3 (47.0, 51.5) |  |
| **Time since diagnosis, yrs** |  |  |  |
| ≤25 | 366 (47) | 49.1 (47.6, 50.6) | 0.092 |
| >25 | 408 (53) | 47.0 (45.6, 48.4) |  |
| **History of relapse** |  |  |  |
| No | 691 (89) | 47.9 (47.0, 48.8) | 0.711 |
| Yes | 83 (11) | 48.4 (45.9, 51.0) |  |
| **Treatment exposures** |  |  |  |
| **Radiation** |  |  |  |
| **Cranial** |  |  |  |
| No | 654 (85) | 48.2 (47.3, 49.1) | 0.247 |
| Yes | 120 (16) | 46.8 (44.5, 49.0) |  |
| **Chest** |  |  |  |
| No | 688 (89) | 47.9 (47.0, 48.8) | 0.748 |
| Yes | 86 (11) | 48.4 (45.6, 51.2) |  |
| **TBI and/or abdominal** |  |  |  |
| No | 707 (91) | 48.0 (47.2, 48.9) | 0.746 |
| Yes | 67 (9) | 47.5 (44.6, 50.5) |  |
| **Glucocorticoids** |  |  |  |
| No | 444 (57) | 47.5 (46.1, 48.9) | 0.420 |
| Yes | 330 (43) | 48.6 (46.8, 50.4) |  |
| **Anthracyclines** |  |  |  |
| No | 482 (62) | 47.6 (46.5, 48.7) | 0.345 |
| Yes | 292 (38) | 48.6 (47.1, 50.0) |  |
| **Alkylating agents** |  |  |  |
| No | 455 (59) | 47.6 (46.5, 48.8) | 0.372 |
| Yes | 319 (41) | 48.5 (47.1, 49.8) |  |
| **Hematopoietic stem cell transplantation** |  |  |  |
| No | 744 (96) | 48.1 (47.2, 48.9) | 0.246 |
| Yes | 30 (4) | 45.5 (41.2, 49.8) |  |

<sup>1</sup> CNS, central nervous system; CCS, childhood cancer survivors; ICC3, *International Classification of Childhood Cancer*, 3<sup>rd</sup> edition; TBI, total body irradiation.
<sup>2</sup> Adjusted for sex, age at survey, and ICC3-3 diagnosis.

## DISCUSSION

### Principal findings

We found that adherence to national dietary intake recommendations and diet quality among long-term CCSs was generally as poor as that of the Swiss population. The diet of CCSs appears particularly low in fiber and too high in saturated fat, though reported alcohol intake was lower in CCSs than in the general adult population. In terms of AHEI, the scores for vegetables and for red and processed meat were better in CCSs than in the general population, whereas scores for sugar-sweetened beverages and nuts, seeds, legumes, and tofu were similarly poor in both groups. Baseline clinical characteristics were not associated with diet quality at follow-up in CCSs. Our findings can inform the promotion of healthy eating in CCSs in Switzerland, who are at increased risk and earlier onset of chronic diseases.

### Strengths and limitations

To assess dietary quality, we calculated the AHEI for CCSs based on FFQ data and for participants in the Swiss National Nutrition Survey based on two 24HDR, which might not have captured all episodically consumed foods. The different dietary assessment tools could have led to different results. MenuCH data are, however, less likely to have led to underreporting, as illustrated by the higher estimated total energy intake (2283 kcal vs 2022 kcal in Bus Santé, 1923 kcal in CoLaus, and 1639 kcal in CCSs). This underreporting should however have little impact on the AHEI. The FFQ is a self-reported questionnaire designed to assess habitual intakes, whereas the 24HDR is an in-depth interview to assess in detail everything a participant has eaten over the past 24 hours. Food products, that are rarely eaten are expected to be less frequently reported in 24HDRs than FFQs. To calculate the dietary intake of CCSs and participants of the Bus Santé and CoLaus study we used the French Centre d’Information sur la Qualité des Aliments food-composition table, whereas we used the Swiss Food Composition Database (SFCDB) for the Swiss National Nutrition Survey. Within the SFCDB, food composition data from branded products relied on information on product packaging or websites provided by manufacturers. This could have resulted in slight dietary differences, although they are expected to be minimal. Differences in dietary intake could have further emerged among study populations because data on pregnancy and lactation were unavailable for the participants of the CoLaus and Bus Santé studies. However, since the Bus Santé study only included women aged 35 years and older, and the CoLaus study women of 40 years and older the proportion of pregnant and lactating women is expected to have been relatively low. Finally, our data were based on a cross-sectional analysis, which precludes the timing of poor dietary intake and quality. Dietary intake of CCSs could change over time due to the occurrence of chronic health conditions or cancer recurrence (46, 47).

Despite these limitations, this is the first study that directly compares dietary intake and quality of CCSs with the general adult population. We included two comparison groups (Bus Santé and CoLaus) from the French speaking region in Switzerland, which assessed diet with the same FFQ as the CCSs. We included one nationwide comparison group (Swiss National Nutrition Survey) that was performed within a similar age range. The SCCSS is strengthened by its national coverage, large sample size, and high response rate, which makes our results representative. Furthermore, we had access to high quality clinical information extracted from the SCCR.

### Adherence to dietary guidelines, diet quality: results in relation to other studies

Our finding that the majority of CCSs does not fully adhere to national dietary recommendations is in line with previous studies among CCSs (6, 10, 14, 15, 48, 49). The low fiber intake we observed, of 12 g/day compared to the recommended DACH intake of ≥30 g/day (43), has also been reported for both short-term and long-term CCSs (6, 10, 48, 49). The St. Jude Lifetime Cohort Study found intakes between 17 and 18 g/day; among 1598 adult CCSs 26 years postdiagnosis (women: 17 g/day, men: 18 g/day) (6) and 2570 adult CCSs 24 years postdiagnosis (17 g/day). Smaller US studies with shorter follow-up times have also found similar results. A fiber intake of 17 g/day was found in 170 CCSs 9 years postdiagnosis (48), but the Chicago Healthy Living Study of 431 adult CCSs 19 years postdiagnosis found a fiber intake of 9 g/1000 kcal compared to the recommended intake of the 2010 Dietary Guidelines for Americans of 14 g/1000kcal (49). The fiber intakes of CCSs seem to be even lower than the intake of the general population (49), which we also observed in our study. This finding is particularly salient for CCSs, who have a higher risk of developing chronic health conditions such as type II diabetes and CVD, and for whom high-fiber foods and whole grains are recommended for prevention and management of these chronic conditions (2, 3). Whether the adherence of CCSs to dietary recommendations is indeed lower than the general population will need to be confirmed in future studies.

Our findings on low overall diet quality are in line with data from the St. Jude Lifetime Cohort of 2570 adult CCSs with a comparable time from diagnosis (24 ± 8 years). They found a mean ± SD Healthy Eating Index 2010 score (HEI-2010) of 57.9 ± 12.4 out of a maximum score of 100 (10). Similar results were observed in other, smaller studies from the US; mean diet quality in CCSs ranged from 33-59% of the maximum score (11–13, 50). Although diet quality in these studies was measured with a different scoring method and is therefore not directly comparable with our results, they all show that diet quality in CCSs is poor. We found that the general population scored even lower, but these lower results may be the result of different dietary assessment tools (FFQ versus 24HDR) or biased reporting, and need to be compared and interpreted with caution. Discrepancies in AHEI component scores between CCSs and the general population were seen particularly for fish intake. Since fish is consumed rarely in Switzerland, the probability of assessing intake on two recall days, as was done in the Swiss National Nutrition Survey, is low (42). Fish consumption may have been more easily assessed in a FFQ, which is used in the SCCSS. This might explain the lower AHEI score in the general population compared to the CCSs.

We did not find differences in AHEI scores between subgroups of CCSs based on clinical characteristics. Compared to a larger study of the St. Jude Lifetime Cohort, we found a similar tendency that survivors of lymphoma had the highest AHEI score (49.6), followed by survivors of leukemia (48.3), whereas survivors of CNS tumors had the lowest score (45.5, *P* = 0.08) (10). Lower diet quality scores among CNS tumor survivors may be the result of the exposure to several risk factors such as cranial radiation therapy (CRT) and surgical damage that can lead to hypothalamic obesity (51, 52). We did not find clear differences between AHEI scores when we looked at treatment exposures. This was consistent with previous literature on exposure with glucocorticoids, anthracyclines, and alkylating agents (10, 12), but inconsistent for abdominal radiation. In the St. Jude Lifetime Cohort, survivors who received higher abdominal radiation doses had lower dietary quality scores compared to those who received lower doses (10). We lacked power to stratify by dose and could not confirm these results. A smaller study including 91 CCSs showed that survivors exposed to CRT had lower HEI scores than those not exposed (12). We found a similar trend, as did a US study among 22 ALL and lymphoma survivors (11). No trend was seen for CRT stratified by dose in the St. Jude Lifetime Cohort among 916 CCSs who received CRT (10). Since cancer diagnosis directly affects the type of treatment a patient receives, occurrence of late effects can play an important role in the survivors dietary eating habits. Large-scale prospective studies with a homogenous study population and repeated dietary assessments are therefore needed to better investigate diet quality between patients exposed to different types of treatment.

### Implications and recommendations

In cancer prevention campaigns, the Swiss Cancer League (www.liguecancer.ch) emphasizes increasing fruit and vegetable consumption and reducing alcohol and red and processed meat intake. This may be based upon AHEI scores for these components that are higher than those of the general population, which are in line with our previous findings (42). However, it is unclear to what extent CCSs are aware of these cancer prevention campaigns, whether dietary recommendations are communicated to them by health care professionals, and whether they perceive diet as a risk factor for late effects. Current CCSs guidelines have general dietary recommendations for all cancer types and treatments combined (9, 53) or do not focus on diet at all (8). However, given the strong evidence concerning diet and health in general and the increasing data for CCSs (5), more focus should be placed on the importance of balanced eating habits and physical activity during annual long-term follow-up visits. The importance of a healthy lifestyle was also confirmed in a study of 1598 CCSs 26 years postdiagnosis (6), which found that an unhealthy lifestyle based on anthropometric, FFQ, and physical activity data according to World Cancer Research Fund/American Institute for Cancer Research recommendations was associated with higher risk of metabolic syndrome. This suggests that CCSs could lower their elevated risk of developing late effects after cancer treatment through behavioral changes. Because CCSs have a dietary intake and quality that is similar to the general population the focus should be on population-based interventions rather than individual counselling.

### Conclusions

We found a poor dietary intake and diet quality in CCSs in Switzerland that was as poor as that reported in the general adult population. Low fiber intake and the high intake of sugar-sweetened beverages and saturated fat are of particular concern. Our findings suggest that previous cancer treatment exposure and cancer characteristics such as age at and time since diagnosis do not substantially influence the diet quality of CCSs in the long term. We suggest population-based interventions reinforcing the importance of a healthy diet to everyone.

Members of the Swiss Pediatric Oncology Group (SPOG) Scientific Committee are as follows: R. Ammann (Bern); R. Angst (Aarau); M. Ansari (Geneva); M. Beck Popovic (Lausanne); P. Brazzola (Bellinzona); J. Greiner (St. Gallen); M. Grotzer (Zurich); H. Hengartner (St. Gallen); T. Kuehne (Basel); K. Leibundgut (Bern); F. Niggli (Zurich); J. Rischewski (Lucerne); N. von der Weid (Basel).

## ACKNOWLEDGEMENTS

The authors express their gratitude to all childhood cancer survivors in Switzerland for filling in the questionnaire and supporting this study. Additionally, we thank the Bus Santé and CoLaus scientific committees for access to their data, and the Swiss Federal Food Safety and Veterinary Office (FSVO), www.blv.admin.ch for providing the Swiss National Nutrition Survey menuCH 2014-2015, Version 2.0 dataset (May 2017). We thank the study team of the Swiss Childhood Cancer Survivor Study (Rahel Kuonen, Erika Brantschen Berclaz, Grit Somer, Annette Weiss, Christina Schindera, Nicolas Waespe, Carole Dupont, Maria Otth, and Cecilia Ferrari), the data managers of the Swiss Paediatric Oncology Group (Claudia Anderegg, Pamela Balestra, Nadine Beusch, Rosa-Emma Garcia, Franziska Hochreutener, Friedgard Julmy, Nadia Lanz, Rodolfo Lo Piccolo, Heike Markiewicz, Annette Reinberg, Renate Siegenthaler, and Verena Stahel), and the team of the Swiss Childhood Cancer Registry (Verena Pfeiffer, Katherina Flandera, Shelagh Redmond, Meltem Altun, Parvinder Singh, Vera Mitter, Elisabeth Kiraly, Marlen Spring, Christina Krenger, and Priska Wölfli). Finally, we would like to thank Christopher Ritter for editorial assistance.

## AUTHORS’ CONTRIBUTIONS & CONFLICT OF INTEREST

The authors’ responsibilities were as follow. FNB conducted the statistical analyses and wrote the manuscript. AC helped with analyzing menuCH data and AHEI scoring. AC, RK, CEK, and MB contributed to the concept and the design of the study, and gave support in the statistical analyses. All authors have revised earlier drafts, and read and approved the final manuscript. None of the authors reports any conflict of interest related to the study.

## Abbreviations

24HDR: 24-hour dietary recall
AHEI: Alternative Healthy Eating Index
ALL: acute lymphoblastic leukemia
BMI: body mass index
CCSs: childhood cancer survivors
CI: confidence interval
CNS: central nervous system
COG-LTFU: Children’s Oncology Group Long-Term Follow-up
DACH: Germany (D), Austria (A), and Switzerland (CH)
CoLaus: Cohorte Lausannoise
CRT: cranial radiation therapy
CVD: cardiovascular disease
Dx: diagnosis
FFQ: food frequency questionnaire
Gy: gray
HEI: Healthy Eating Index
HSCT: hematopoietic stem cell transplantation
ICCC-3: International Classification of Childhood Cancer, 3^rd^ edition
IQR: interquartile range
OR: odds ratio
PAFQ: physical activity frequency questionnaire
SCCR: Swiss Childhood Cancer Registry
SCCSS: Swiss Childhood Cancer Survivor Study
SFCDB: Swiss Food Composition Database
SHS: Swiss Health Survey
UK: United Kingdom
US: United States

## Notes

**Sources of support:**This study is supported by Swiss Cancer Research (KLS-3644-02-2015). The work of the Swiss Childhood Cancer Registry is supported by the Swiss Pediatric Oncology Group (www.spog.ch), Schweizerische Konferenz der kantonalen Gesundheitsdirektorinnen und–direktoren (www.gdk-cds.ch), Swiss Cancer Research (www.krebsforschung.ch), Kinderkrebshilfe Schweiz (www.kinderkrebshilfe.ch), the Federal Office of Health (FOH), and the National Institute of Cancer Epidemiology and Registration (www.nicer.org).

