## Supplementary material for "Dietary intake and diet quality of Swiss adult survivors of childhood cancer compared to the general population"

### ONLINE SUPPLEMENTAL MATERIAL (OSM)

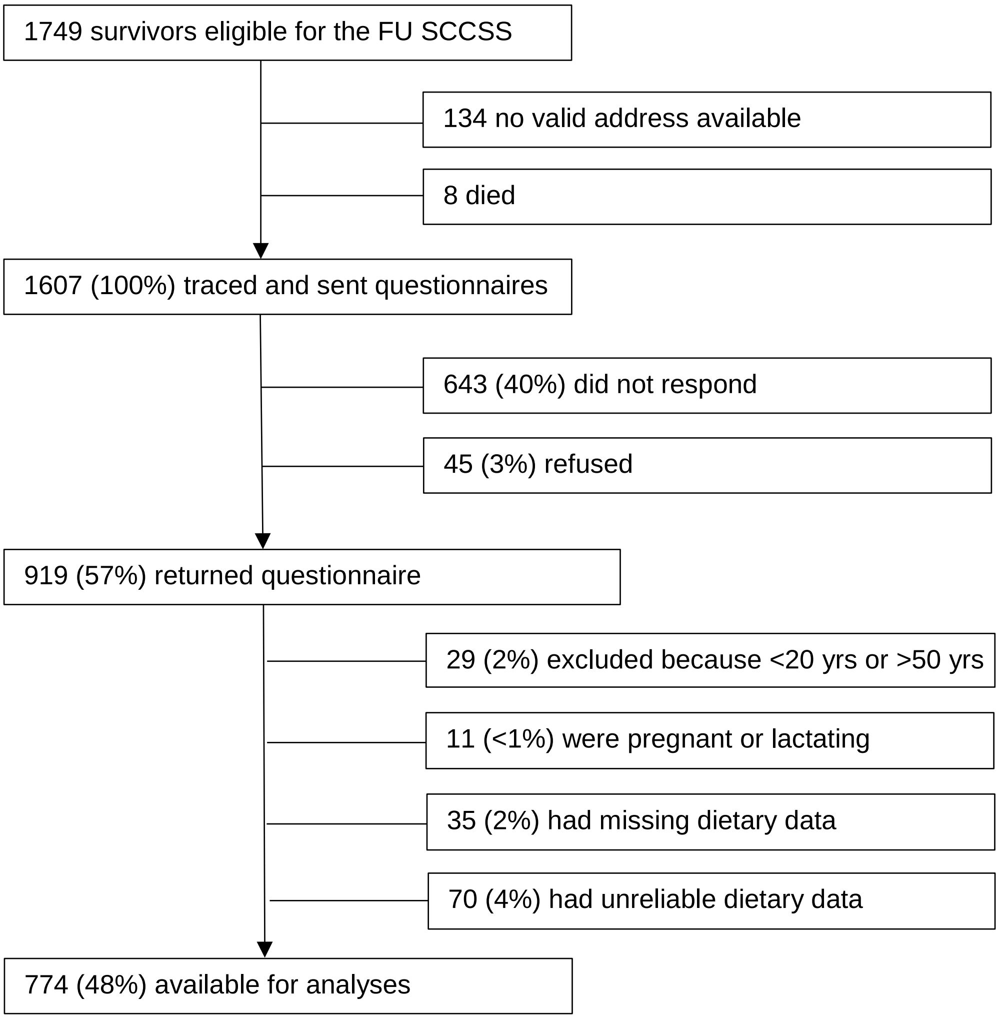


**Figure S1** Response rates in the follow-up Swiss Childhood Cancer Survivor Study[1]

[1] FU, follow-up; SCCSS, Swiss Childhood Cancer Survivor Study.

**Table S1** Clinical characteristics of childhood cancer survivors[1]

| Characteristics | CCSs (*n* = 774) |
|---|---|
| **ICCC3 diagnosis**, *n (%)* | |
| I: Leukemia | 237 (31) |
| II: Lymphoma | 163 (21) |
| III: CNS tumor | 79 (10) |
| IV: Neuroblastoma | 28 (4) |
| V: Retinoblastoma | 12 (2) |
| VI: Renal tumor | 52 (7) |
| VII: Hepatic tumor | 6 (1) |
| VIII: Bone tumor | 49 (6) |
| IX: Soft tissue sarcoma | 64 (8) |
| X: Germ cell tumor | 40 (5) |
| XI & XII: Other tumor | 25 (3) |
| Langerhans cell histiocytosis | 19 (2) |
| **Age at diagnosis**, *n (%)* | |
| <5 yrs | 251 (32) |
| 5-9 yrs | 163 (21) |
| 10-14 yrs | 224 (29) |
| 15-20 yrs | 136 (18) |
| **Time since diagnosis**,[2] yrs | 25.7 (20.1-31.1) |
| **History of relapse**, *n (%)* | 83 (11) |
| **Treatment exposures**, *n (%)* | |
| **Radiation**, *n (%)* | |
| Any | 276 (36) |
| Cranial | 120 (16) |
| <18 Gy | 27 (3) |
| ≥18 Gy | 93 (12) |
| Chest | 86 (11) |
| <30 Gy | 33 (4) |
| ≥30 Gy | 53 (7) |
| TBI and/or abdominal | 67 (9) |
| **Glucocorticoids** | 330 (43) |
| **Anthracyclines** | 292 (38) |
| **Alkylating agents** | 319 (41) |
| **Hematopoietic stem cell transplantation** | 30 (4) |

[1] CCS, childhood cancer survivor; CNS, central nervous system; Gy, gray; ICCC3, *International Childhood Cancer Classification,* 3rd edition; TBI, total body irradiation.

[2] Values are medians (IQRs).

**Table S2** Dietary intake in childhood cancer survivors by sex compared to the DACH dietary recommendations[1]

| Nutrients | DACH recommendations[2] | | CCSs (*n* = 774) | | | |
|---|---|---|---|---|---|---|
| | men | women | men (*n* = 386) | | women (*n* = 388) | |
| | | | mean ± SD | % DACH[4] | mean ± SD | % DACH[4] |
| Total energy, kcal[3] | 2300-3400 | 1800-2600 | 1743 ± 632 | 61 | 1535 ± 508 | 70 |
| **Macronutrients** | | | | | | |
| Protein, g | 57 | 48 | 73.8 ± 33.8 | 129 | 58.5 ± 22.1 | 122 |
| % of energy[5] | 10-20 | 10-20 | 16.9 ± 3.7 | 112 | 15.4 ± 3.4 | 103 |
| Vegetal, g | | | 18.3 ± 7.7 | | 18.0 ± 7.5 | |
| Animal, g | | | 55.5 ± 30.7 | | 40.6 ± 19.9 | |
| Carbohydrates, g[6] | 225-275 | 225-275 | 187 ± 76.2 | 75 | 174 ± 71 | 70 |
| % of energy[5, 6] | 45-55 | 45-55 | 42.8 ± 8.4 | 86 | 45.2 ± 9.1 | 90 |
| Mono/disaccharides, g | | | 87 ± 46 | | 84 ± 46 | |
| Polysaccharides, g | | | 98 ± 46 | | 90 ± 42 | |
| Total fiber, g[6] | ≥30 | ≥30 | 11.5 ± 5.8 | 38 | 12.9 ± 6.9 | 43 |
| Total fat, g | | | 72.2 ± 30.2 | | 64.2 ± 25.4 | |
| Total fat, % of energy[5] | 30-40 | 30-40 | 37.2 ± 6.6 | 106 | 37.5 ± 7.2 | 107 |
| Saturated fat, g | | | 27.3 ± 12.5 | | 23.1 ± 10.4 | |
| Saturated fat, % of energy[7] | <10 | <10 | 14.0 ± 2.9 | 140 | 13.5 ± 3.4 | 135 |
| Monounsaturated fat, g | | | 29.9 ± 13.3 | | 27.3 ± 11.9 | |
| Polyunsaturated fat, g | | | 9.0 ± 3.9 | | 8.2 ± 3.6 | |
| Cholesterol, mg[5] | 300 | 300 | 376 ± 209 | 125 | 310 ± 202 | 103 |
| **Micronutrients** | | | | | | |
| ***Vitamins*** | | | | | | |
| A, mg-RE[8] | 1.0 | 0.8 | 0.7 ± 0.7 | 71 | 0.7 ± 0.7 | 89 |
| D, µg | 20 | 20 | 2.5 ± 2.0 | 12 | 2.3 ± 2.0 | 12 |
| ***Minerals*** | | | | | | |
| Calcium, mg | 1000 | 1000 | 908 ± 502 | 91 | 869 ± 437 | 87 |
| Iron, mg | 10[4] | 15 | 9.7 ± 3.9 | 97 | 8.3 ± 3.1 | 56 |
| Alcohol, g[9] | 20 | 10 | 7.5 ± 7.9 | 37 | 4.0 ± 7.2 | 40 |

[1] CCS, childhood cancer survivors, DACH, Dietary recommendations for Germany (D), Austria (A) and Switzerland (CH).
[2] DACH recommendations for the general population age 20-50 yrs, excluding pregnant and lactating women, 2015.
[3] Depending on physical activity level (PAL, 1.4-1.8) and age.
[4] Percentage of mean intake to the DACH recommended intake level * 100. Recommended intake is estimated on the basis of the age-sex groups of the DACH guidelines, weighted by the age and sex distribution of the study population. For alcohol intake the maximum tolerated dosage was taken.
[5] Federal Food Safety and Veterinary Office FSVO, www.naehrwertdaten.ch.
[6] BLV (2009). Kohlenhydrate in unserer Ernährung – Empfehlungen des BLV. www.blv.admin.ch.

[7] BLV (2012). Fett in unserer Ernährung – Empfehlung des BLV. www.blv.admin.chv, recommendations as a percentage of the daily energy demand and based on a daily energy intake of 2000 kcal.
[8] Total vitamin A= retinol + carotene/12, expressed in µg RAE.
[9] Maximum tolerated dosage.

**Table S3** Scoring method and mean scores (95%CI) of components of the modified Alternate Healthy Eating Index[1,2] in childhood cancer survivors and the general population (menu.CH) by sex

| Components | Criteria for minimum score (0) | Criteria for maximum score (10) | Score in CCSs[12] mean (95%CI) | | Score in Menu.CH[13] mean (95%CI) | |
|---|---|---|---|---|---|---|
| | | | men ($n$ = 386) | women ($n$ = 388) | men ($n$ = 498) | women ($n$ = 636) |
| Vegetables, excluding potatoes (servings/day)[3] | 0 | ≥ 5 | 4.6 (4.4, 4.9) | 5.5 (5.3, 5.7) | 3.2 (3.2, 3.2) | 3.3 (3.3, 3.3) |
| Fruit, excluding juice (servings/day)[4] | 0 | ≥ 4 | 2.9 (2.6, 3.2) | 3.9 (3.6, 4.2) | 2.9 (2.9, 2.9) | 3.4 (3.4, 3.5) |
| Whole grains (g/day)[5] | | | | | | |
| Men | 0 | ≥ 90 | 3.3 (3.0, 3.6) | | 3.4 (3.4, 3.4) | |
| Women | 0 | ≥ 75 | | 4.7 (4.4, 5.0) | | 4.0 (4.0, 4.0) |
| Sugar-sweetened beverage and fruit juice (servings/day)[6] | ≥ 1 | 0 | 3.8 (3.4, 4.2) | 4.9 (4.5, 5.3) | 3.8 (3.7, 3.9) | 4.8 (4.7, 4.9) |
| Nuts, seeds, legumes, and tofu (servings/day)[7] | 0 | ≥ 1 | 1.9 (1.6, 2.2) | 2.6 (2.3, 2.9) | 1.7 (1.6, 1.7) | 2.5 (2.5, 2.6) |
| Red and processed meat (servings/day)[8] | ≥ 1.5 | 0 | 3.5 (3.2, 3.8) | 5.8 (5.5, 6.1) | 2.3 (2.3, 2.4) | 5.2 (5.2, 5.2) |
| Trans fat (% of total energy intake)[9] | ≥ 4 | ≤ 0.5 | 9.6 (9.5, 9.7) | 9.5 (9.5, 9.6) | 9.5 (9.5, 9.5) | 9.4 (9.4, 9.4) |
| Fish, excluding processed products (g/day) | 0 | 32.4 | 5.2 (4.8, 5.5) | 4.9 (4.6, 5.3) | 2.6 (2.6, 2.6) | 2.6 (2.6, 2.6) |
| PUFA (% of total energy intake)[10] | ≤2 | ≥10 | 4.8 (4.6, 5.1) | 5.2 (4.9, 5.5) | 4.4 (4.3, 4.4) | 4.3 (4.3, 4.3) |
| Alcohol (drinks/day)[11] | | | | | | |
| Men | ≥ 3.5 | 0.5 - 2.0 | 5.4 (5.0, 5.7) | | 5.0 (4.9, 5.0) | |
| Women | ≥ 2.5 | 0.5 - 1.5 | | 3.9 (3.6, 4.3) | | 4.0 (4.0, 4.1) |
| **Total** | **0** | **100** | **45.0 (43.8, 46.2)** | **50.9 (49.7, 52.1)** | **38.7 (38.4, 38.9)** | **43.8 (43.6, 44.0)** |

[1] Adapted from Chiuve et al. J Nutr 2012 142(6):1009-18
[2] Intermediate food intake was scored proportionately between the minimum score 0 and the maximum score 10.
[3] One serving was equal to 118.3g of raw or cooked vegetables or 250g of vegetable soup. Dried vegetables were not included.
[4] One serving was equal to 118.3g of raw or cooked fruit. Dried fruit was not included.
[5] Whole grain products like whole grain bread and muesli were included.
[6] One serving was equal to 226.8g.
[7] One serving was equal to 28.4g.
[8] One serving was equal to 113.4g of red meat or 42.5g of processed meat.
[9] Consumption was estimated based on the assumption that each food contains a maximum of 2g of trans fat per 100g of total fat, as defined in Swiss regulation.
[10] The highest score was given to individuals with ≥10% of total energy intake from polyunsaturated fatty acids (PUFA).
[11] One drink was 113.4g of wine, 340.2g of beer or 42.5g of liquor. A score of 2.5 was given to nondrinkers.
[12] Adjusted for age at survey and ICCC-3 diagnosis
[13] Adjusted for age at survey and standardized on sex, age at survey, and language region according to CCSs.
